# Modelling Plant Cortical Microtubules: Curvature Sensing from Bending Energy

**DOI:** 10.64898/2026.06.22.733854

**Authors:** Tim Y.Y. Tian, Geoffrey Wasteneys, Colin B. Macdonald, Eric N. Cytrynbaum

## Abstract

Within plant cells, the self-organization of cortical microtubules (MTs) into ordered arrays is an important process for directional growth. There is growing evidence that cortical MTs respond to cell shape and/or mechanical stresses in the cell wall, requiring *in silico* models on complicated surfaces to provide a complete understanding. Most models assume that MTs are directionally persistent, following geodesics of the surface. This ignores the expected tendency of these elastic filaments to minimize curvature. Our recent model incorporated minimization of MT curvature in cylindrical cells and found curvature to be significant in biasing the array organization. Here, we generalize to a larger class of surfaces, modelling individual microtubule shapes to provide insights into the role of geometric cues and highlight differences with previous models that use the geodesic assumption. We first show that geodesic models, including current models with finite persistence lengths, exhibit an invariance across certain geometries, leading to biophysically counterintuitive results. Incorporating curvature minimization, we show the difficulties imposed by high-curvature cell edges, elucidating potential new roles of proteins in helping microtubules traverse edges. Lastly, we show that geometries with competing curvature cues result in diverse curves previously not considered. These results provide geometric intuition for how various cell geometries affect individual cortical microtubules, helping us to better understand the processes required for the establishment of microtubule arrays in broad contexts such as: bundles spanning adjacent cell faces in prism-like root and leaf epidermis cells, protruding geometries of trichome cells, and rounded surfaces such as confined protoplasts.

## 1 Introduction

Plant cells must coordinate the orientation of the cellulose microfibrils in the cell wall that provide reinforcement against mechanical stresses. Evidence suggests that this coordination is facilitated by an array of ordered and similarly oriented microtubules (MTs) at the cell cortex. In the case of elongating cells in organs such as roots and stems, MT arrays and cellulose microfibrils are oriented transverse to the axis of elongation to prevent radial expansion. In addition, MTs appear in various configurations within other cell types at different stages of development. Examples are shown in Fig. 1A: *Arabidopsis* (i) prism-like root division-zone cells [2], (ii) protrusion-shaped trichome cells [44], and (iii) ellipsoidal protoplasts [12]. The mechanisms that orient MTs is an active area of inquiry [38, 12]. It remains unclear what physical properties MTs are responding to: candidates include mechanical stress [12], strain, cellulose feedback [46], and regulation by MT-associated proteins (see [50, 2, 41, 29, 15], among others). Furthermore, it is unclear how MTs would respond to these properties; what biological pathways are responsible for propagating such signals? Among these ideas is the prominent hypothesis that MTs sense mechanical stress. Hamant et al. [22] suggest that the rigidity of MTs enable these polymers to maintain their direction over the length scale of a cell, making them well-suited to sense cell-scale mechanical signals.

**Figure 1.**
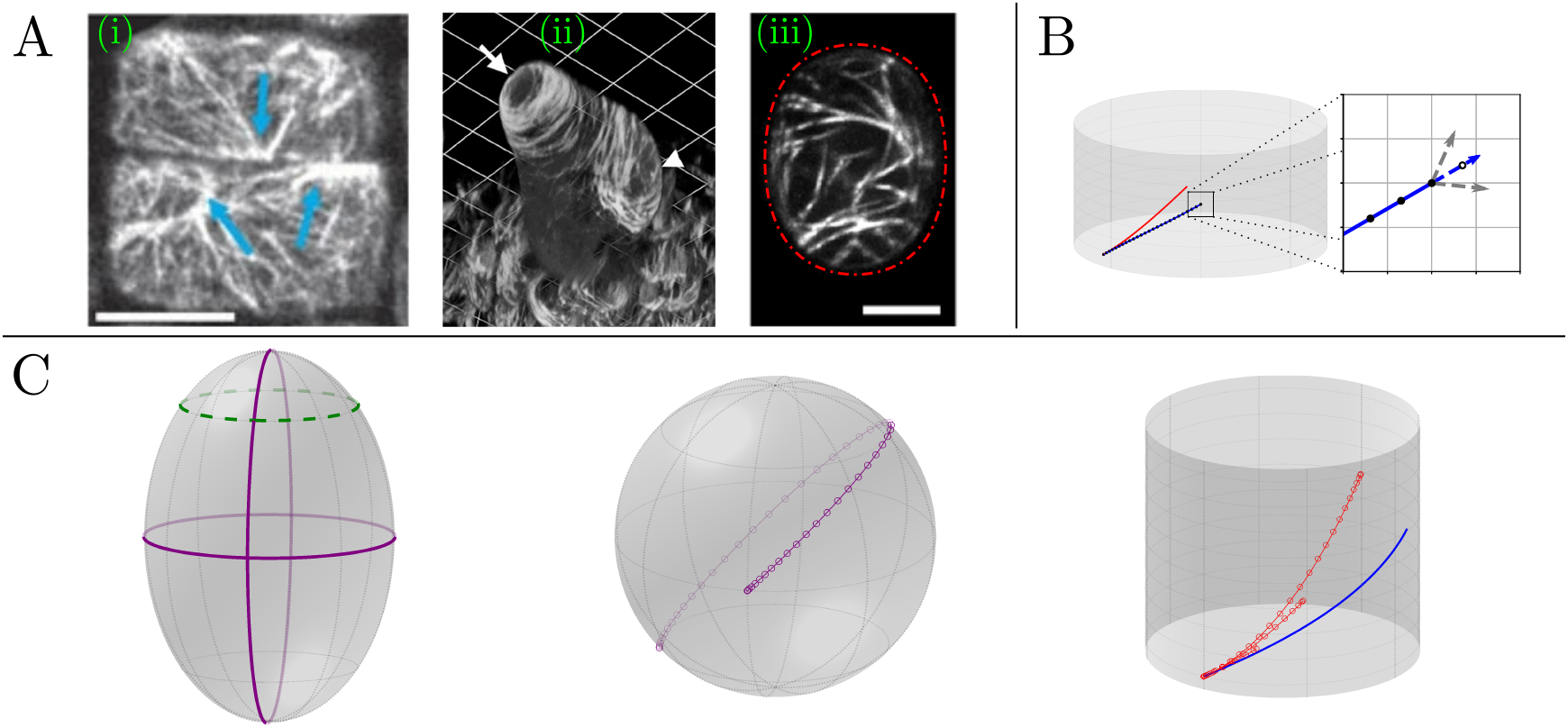
Cortical microtubules in various geometries and illustrations of elastic curves and geodesics on various surfaces. A: Examples of cortical microtubule arrays (white) in *Arabidopsis* (i) root division zone epidermal cells (image from [2, Fig. 4a]), (ii) trichomes (image from [44, Fig. 2E]), and (iii) protoplasts (image from [12, Fig. 2E], red outline added). Arrows indicate (i) bundles of MTs traversing sharp edges between two cell faces and (ii) MT-depleted regions at trichome tips. Scale bars in (i) and (iii) are 5*µ*m and a unit grid in (ii) is 7.31*µ*m. B: An elastic curve (red) and geodesic (blue) of the same length and initial direction are plotted on a cylinder [45]. The entire elastic curve consists of a free MT tip. The geodesic is composed of an infinite number of anchored sections (anchors are shown in the inset as finitely spaced black dots for illustration only). B inset: A MT with a short free tip length avoids the energetic cost of bending laterally (gray arrows) toward a lower curvature direction because it is too short to benefit enough from such a redirection. Hence, it follows a geodesic. C: Elastic and non-elastic curves on various surfaces. Left: a spheroid. This is an example of a surface of revolution, for which elastic curves have a well-developed theory [36]: curves along any line of longitude (vertical purple ellipse) extremize the bending energy; curves along circles of latitude (e.g. dashed green) generally do not extremize the curvature energy, except if the circle of latitude is also geodesic (horizontal purple ellipse). Middle and right: A comparison of elastic curves and geodesics on a sphere and cylinder, both of which have existing methods for calculating elastic curves [36]. On the sphere, the solid purple curve is part of a great circle which is both an elastic curve and a geodesic; the same result is calculated using our energy minimization method (purple dots, see Sec. 4.2). On the cylinder, the elastic curves are calculated using our energy minimization (red dots) and Euler-Lagrange equations (red curve) [5].

Understanding the correlation between MT orientation and cell shape or changes in cell shape has motivated studies of more complex geometries in both experiments and theoretical models. On the experimental side, there have been *in vitro* studies of protoplasts confined to geometries of various shapes and aspect ratios, such as ellipsoids and rounded prisms [12, 14]. In general, it appears that MTs in protoplasts with geometric asymmetries are biased toward the longitudinal axis by default [12, 14]. Notably, MT orientation responds to changes in pressure, suggesting that tensile stress in the membrane is the signal [12]. On the modelling side, simulations have been generalized to complicated geometries [10, 11, 35, 14]. Often, these studies invoke the modification of MT dynamics on cell faces or edge-induced catastrophes between cell faces to reproduce experimental observations. However, these models do not incorporate the bending mechanics of MTs on curved surfaces — MTs are assumed to travel along geodesics (discussed further below). The omission of curvature minimization by these rigid filaments means that our understanding of the effects of geometry on MT arrays remains incomplete. This incompleteness is especially concerning with regard to studies of Nitella internodal cells [37, 52] and confined protoplasts, where there are no clear cell faces or edges to modify MT dynamics at different regions of the cell (or where the edges are far apart in the case of Nitella internodal cells).

To understand the biological processes at play, it is necessary to first understand the behaviour of MTs in the context of cellular geometry alone, in the absence of additional layers of regulation. Can geometry and microtubule mechanics explain the observed MT organization?

If not, do they contribute to or antagonize the organization process? *In vitro* experiments on persistence lengths [18, 40] suggest MTs are relatively stiff polymers and should minimize their bending. In the case of a MT in free space, or constrained to a plane, the preferred MT configuration is straight. In the case of MTs constrained to a curved surface, MTs should assume low curvature shapes in the absence of any other biological processes. This is consistent with *in vitro* experiments on MT coiling in confined geometries [27] and suggests a general principle: MTs ought to follow regions of minimal curvature on the surface. In the simple case of a cylinder, there is a uniform influence of curvature — the transverse direction induces curvature in MTs whereas the longitudinal direction does not. Therefore, MTs should align preferentially in the longitudinal direction. The strength of this influence depends on how much the MTs sense the curvature. The longer the unanchored section of the MT at the growing plus end (referred to here as the “free tip”) is prior to anchoring, the more the MT tends toward the longitudinal direction [5]. For short free tips, bending laterally is more energetically costly than the energetic benefit of redirecting to the low curvature direction, which is why MTs follow geodesics in the limit of a high anchoring rate [5]. We recently showed that, in cylindrical cells, the influence of curvature should extend beyond single MTs to bias the entire MT array toward the longitudinal direction [46]. This is not observed *in vivo* and it is necessary to invoke some other active biological processes to explain the observed transverse arrays. Here we ask: how does the influence of curvature generalize to more complex geometries, such as ellipsoids, which serve as a model for confined protoplasts? To answer this question, it is necessary to address existing misconceptions and develop a deeper understanding of curvature-minimizing MTs on curved surfaces.

For a free MT on a two-dimensional plane, MT rigidity results in “directional persistence”: a tendency to be straight due to curvature minimization. In other words, the energy minimizers and straight lines are equivalent. This equivalence no longer holds on curved surfaces in general; energy minimization and directional persistence are different. The former refers to curves extremizing their curvature energy, which we refer to as *elastic curves*. The latter refers to curves that seek to maintain their tangential direction on a surface, known as geodesics.

Notably, geodesics are not curvature-energy minimizers in general. Manning et al. [36], note a long-standing confusion between elastic curves and geodesics even among prominent mathematicians: “Hilbert and Cohn-Vossen [23] incorrectly suggested a flexible knitting needle, constrained to conform to a surface, as one model for a geodesic on a surface. This model actually gives a relaxed elastic line [curve] on the surface, and is not generally a geodesic….” The difference between elastic curves and geodesics is illustrated in Fig. 1B, where an elastic curve (red) and geodesic (blue) of the same length and initial direction are found on a cylinder. For a long elastic curve, corresponding to a free MT tip with a non-zero length from the most recent anchor (leftmost filled black circle) to the plus end, it is energetically beneficial to accept the cost of bending away from its initial tangent direction if that allows it to reach a region of the surface with lower curvature. In the limit of frequent anchoring, the free MT tip gets less benefit from early bending from its initial tangent direction (grey dashed arrows) because there is less length over which to reap the benefits of lower curvature closer to the plus end. In the limit that the distance from the most recent anchor to the plus end goes to zero (infinite anchoring rate), the elastic curve adheres more and more closely to its initial tangent direction (“directional persistence”) eventually approaching a geodesic [5]. With successive growth and anchoring (e.g. empty black circle), in this limit, the entire MT traces out a geodesic (blue).

From a modelling perspective, the incorporation of cell geometry increases the complexity in the already rich mathematical theory of elastic curves. Recent models have examined free-moving (yet surface constrained) filaments in static geometries such as packing infinite filaments into ellipsoids [19], and Langevin dynamics simulations of self-propelling active elastic filaments [26]. There are also models coupling elastic filaments with deformable membranes [7, 42] but the confinement of CMTs to the cortex via anchoring presents a unique context: the filament is typically assumed to be fixed in place by anchors, while being constrained to a rigid shell due to turgor pressure against the cell wall.

In this theoretical context of elastic curves constrained to a surface, Manning et al. [36] derived mathematical conditions for elastic curves and, from their general results, analyzed examples of elastic curves on specific surfaces. They found that the only surfaces on which geodesics and elastic curves are the same are spheres and planes. On planes, these are straight lines and, on spheres, they are great circles, with the spherical case shown in Fig. 1D. For surfaces created by rotating a curve around an axis (a surface of rotation), there are special cases in which geodesics and elastic curves coincide: any arc along a meridian (the intersection between the surface and a plane passing through the axis of rotation) is always both a geodesic and an elastic curve; any arc along a circle of latitude (the locus of a single point on the curve throughout the rotation) that is a geodesic is also an elastic curve, and vice versa. In the case of an ellipsoid of revolution (illustrated in Fig. 1D), the only case of a circle of latitude being a geodesic is the equator. On a circular cylinder, the meridians correspond to any vertical line, and all circles of latitudes are geodesic — therefore any arc along these curves are elastic curves. While it is possible to say that such curves are extrema of the functional Eq. 1, it is not the case that these are all minima. This was shown in [5], where transverse curves become unstable (local maxima) and buckle when the curve is beyond a critical length. Furthermore, in the case of the cylinder, there are previously existing methods to solve for elastic curves corresponding to any initial condition, illustrated in the left panel of Fig. 1D. Additional theoretical results exist in the context of closed curves and/or curves incorporating additional energy terms such as torsion, typically applied to simple geometries such as spheres (see, e.g., [28, 49, 20, 21, 24]). Recent work has examined elastic curves in the specialized context of compressive buckling on quadratic surfaces [53].

Outside of these results, to our knowledge, solving the variational problems to obtain an explicit parametrization of elastic curves remained challenging for general surfaces. Until now, this has been a technical roadblock in developing realistic models of CMT organization in complex geometries. Here, we present a method of numerically solving for elastic curves on smooth surfaces and apply that method to general geometries of biological relevance. We show that the extended lengths of MTs, in combination with their rigidity, makes them curvature sensors. The curvature sensing is non-local in that the MT shape results from the curvature sensed along its entire length rather than at its very tip.

## 2 Results

To understand the non-locality of MT curvature sensing, we studied progressively more complicated shapes by adding compounding curvature influences. As a point of reference, these curves were compared to their geodesic counterparts. For this reason, we start by characterizing geodesics on more complicated surfaces. This gives insight into what aspects of cell geometry the geodesic models are able to sense, and what aspects they miss.

Following that, we present analysis of MTs modelled as elastic curves. MTs were assumed to be purely cortical, firmly anchored at the point of nucleation — mathematically, the curves were constrained to the 2D surface and had a fixed initial position and tangential direction. Other than being on the surface, the elastic curves had no constraints on the free tip, meaning that MTs were anchored only at their nucleation points. This resulted in the curvature minimization problem described in Eq. (1) of the Methods section below.

### 2.1 Invariance of Geodesic Models Across Isometric Surfaces

In the cases of flat planes and circular cylinders, geodesics are straight lines and helices respectively. Adding one layer of complexity, we next looked at generalized cylinders, each resulting from the translation of a planar curve. Examples are shown in Fig. 2. We included several lines of latitude (horizontal lines of constant height) in all figures as a visual aid (darker grey). On the generalized cylinders (Fig. 2A-C), the curvature along lines of latitude was not constant, in contrast with flat planes and circular cylinders. Despite this, geodesics generated curves of constant pitch, just as they were on the simpler surfaces — the angle between the tangent to the geodesic and the line of latitude at any point on the geodesic was the same — suggesting that geodesics are not sensitive to the curvature of the surface. As a result, geodesics on different generalized cylinders with the same initial pitch had the same heights at corresponding points along their lengths. In Fig. 2, the solid and dotted blue curves had the same initial pitch across each surface, and had the same heights across each surface (seen by comparing their heights to the indicated line of latitude). This resulted in two more properties: the distances travelled by the corresponding geodesics on each surface were the same, and the angles between intersecting curves were the same. Together, this means that the two geodesics collided at the same distance from their starting points and had the same angle of collision on each surface.

**Figure 2.**
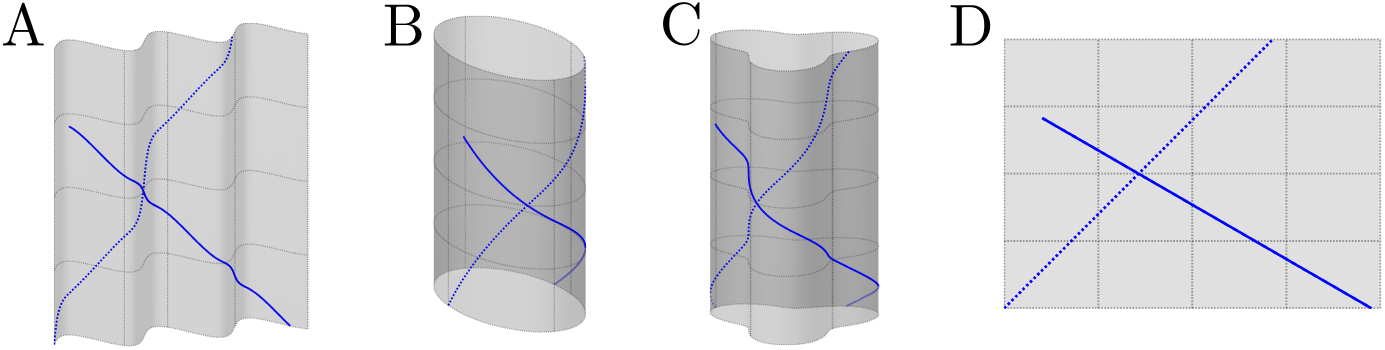
Geodesic curves (solid and dotted blue) as models of CMTs along various surfaces. Each surface is a generalized cylinder, obtained by translating a smooth planar curve: sinusoid (A), oval (B), closed curve with protrusions (C), and straight line (D). For all surfaces shown, each solid blue geodesic has the same initial angle with respect to the horizontal; similarly for the dashed geodesics. The surface-constrained distance between starting points of the solid and dashed curves is the same on all surfaces. The horizontal grid lines have the same spacing on each surface; the vertical grid lines do not.

The invariance of geodesic curves across these surfaces is not a coincidence. Mathematically, this is a consequence of the fact that the plane (Fig. 2D) can be mapped to any smooth generalized cylinder with the lengths of any curves on the plane preserved by the mapping; that is, there is an isometry between a plane and a generalized cylinder. In addition to preserving distances, angles and geodesics are also preserved under an isometry. This means that a model of geodesic CMTs on any smooth generalized cylinder is equivalent to a model on a plane, high-lighting the fact that the direction in which a geodesic MT grows is not affected by the local surface curvature at the growing tip.

That geodesic models on arbitrarily curved cylinders are each equivalent to a model on a plane is contrary to biophysical intuition: one expects MTs to be affected by curvature due to their rigidity. We address the influence of curvature on MT trajectories in the next two sections by considering elastic MTs. In the subsequent sections, figures comparing elastic curves with geodesics contain a MT comparison group described in Fig. 3.

**Figure 3.**
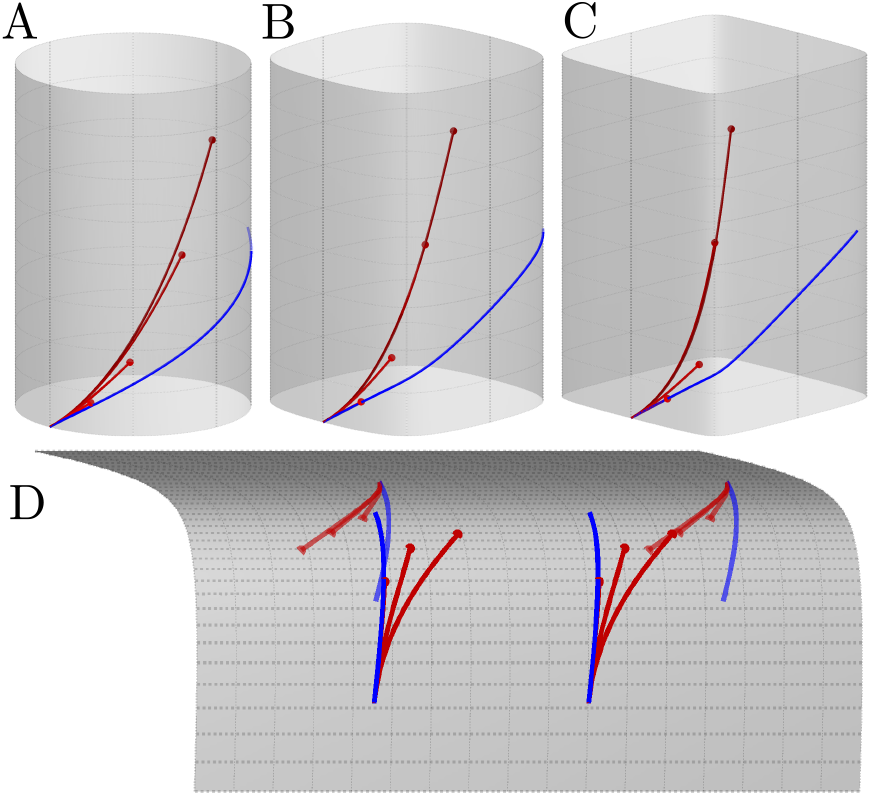
Elastic and geodesic curves on cylinders. Each figure contains a MT comparison group: a set of red elastic curves of various free tip lengths sharing a common initial position and angle, paired with a single blue geodesic with the same initial conditions and the length of the longest elastic curve in the set. A-C: elastic curves on cylinders with progressively sharper edges. Each cylinder has a “half-width” of 1, measured from the centerline of the cylinder to the base of the MTs. The cylinders are interpolations between a circular cylinder and a square cylinder (called “squircular cylinders”) as explained further in the Appendix Fig. 7. The red elastic curves on each squircular cylinder have lengths 0.5, 1, 2, and 3. The tips of the MTs are represented by the dots. D: potential mechanisms for the establishment of transfacial bundles (bundles spanning adjacent cell faces) across sharp edges of squircular cylinder as shown in C, but rotated sideways. Left: two MT groups are illustrated, one group passing from the transverse (top shaded) face to the periclinal face (facing out of the page), and the other passing from the periclinal to the transverse face. Right: the same scenario, with the MTs starting offset so as to align when long enough to deflect in a low anchor density scenerio. When the prism is scaled to have a half-width of 5*µ*m, the minimum radius of curvature in the prism (located at each edge) is 0.94*µ*m, consistent with imaging showing sharp edges to have a radius of curvature of around 1*µ*m [2].

### 2.2 Curvature Sensing from Circular to Square Cylinders

We first compared the curvature effects on MT trajectories through a progression of cylinders from purely circular through to those with sharper edges (i.e., intersections of cell faces), tending to a square-shaped cylinder (a square prism with infinite height). The prism-like shapes serve as more realistic models for cells packed into multicellular organs, such as roots (Fig. 1A(i)) and hypocotyls. In all cases, the influence of curvature was clear: curves deflected longitudinally, as previously shown[27, 5, 45]. MT deflection was non-local in that it depended on the curvature along its entire free length due to the integral in Eq. (1). As shown in Fig. 3A, with the uniform surface curvature of a circular cylinder, MT length was the main factor in how much the MT deflected. On generalized cylinders with varying curvature in their cross-section, curvature is not uniform (Figs. 3, panels B & C).

On the right-most cylinder Fig. 3C with the most square-like shape, the MT initially deflects vertically the least (compared with the other shapes) because its body is in a region of lower curvature. However, as the MT grows into and past the region of high curvature, it deflects more than in the circular cylinder in panel A (as seen by comparing the heights of the MT tips relative the grid lines in each cylinder). This greater overall deflection is a result of the squared dependence on curvature in the energy (Eq. (1)).

This shows that, even if a cell has mostly flat sides, an edge should induce a deflection that, depending on its sharpness, is greater than that of a similar MT on a circularly cylindrical cell. This observation suggests a strategy to mitigate MT deflection at edges: the non-locality of curvature sensing can be modulated by the length of the MT tip via anchoring by proteins such as CLASP (in addition to its role in MT stabilization). This idea is discussed further in Discussion Sec. 3.

The ability of a MT to traverse edges and establish transfacial bundles (bundles spanning adjacent cell faces shown in Fig. 1A(i)) in recently-divided cells is associated with proper cell growth and proliferation in *Arabidopsis* root cells and interdigitation of leaf epidermal cells [3]. This was shown in a previous study [2] that found, in mutants devoid of any CLASP protein, there was both an inability to establish transfacial MT bundles across sharp transverse edges, and a greatly reduced rate of cell proliferation.. In the same study, it was further observed that MTs traversing faces within wild-type cells tended to merge with existing transfacial MTs. In another study [32], mutants lacking in MAP65 displayed similar reductions in root cell proliferation. MAP65s and their animal homologue PRC1 specifically promote the formation of antiparallel microtubules [17, 43].

This motivates a potential alternative mechanism for transfacial bundle creation: MAP65 is necessary for the establishment and persistence of transfacial bundles. The deflection of MTs across a sharp edge could cause antiparallel MTs to deflect toward each other, and MAP65 could then promote their crosslinking to establish the first pioneer bundle. These two possible scenarios — transfacial bundle creation through anchoring or MAP65 — are shown in Fig. 3D. The MT diagram on the left shows that incoming elastic MTs (red) from different faces may deflect away from one another. In such a scenario, it would be beneficial for MTs to have high anchoring while also requiring CLASP to stabilize MTs in the region of slow polymerization due to high curvature. The MT diagram on the right shows an alternative scenario in which the deflection of elastic MTs pushes MTs toward each other — a scenario requiring less of both stabilization and anchoring. Upon establishment of the transfacial bundles, further entrainment of MTs via MAP65 binding would allow for the persistence of the bundles.

### 2.3 Elastic Curves on Geometries with Competing Curvature Cues

The curvature influence on more general geometries is complicated when the surfaces have fewer symmetries than cylindrical surfaces. We studied elastic curves on a paraboloid of revolution as an example of a surface with more spatially heterogeneous regions of high and low curvature compared to cylinders. This surface served as a generic model of cellular protrusions, as found, for example, in trichome cells (Fig. 1A(ii)). For this case, we defined “transverse” as running along or close to circles of latitude, some of which are included as light grey circular rings in Fig. 4. We considered three groups of MTs growing on the paraboloid, shown in Fig. 4. The first group consisted of MTs with one end anchored on a circle of latitude and its direction clamped tangent to the circle of latitude (A). As we expected from theoretical results [36], circles of latitude generally did not minimize curvature. Instead, the MTs deflected downward in the direction of lowest curvature. The avoidance of the cell tip was reminiscent of the MT-depleted zone in trichome cells. If the MTs were clamped at a slight angle upward instead, the direction of lowest curvature was upward for relatively short MTs (B). However, once the MTs were long enough to reach the area around the paraboloid vertex, the energy minimizer quickly became a downward trending curve. This again illustrated the non-locality of MT curvature sensing: the shape of the MT depended on both the length of the unanchored portion of the MT’s tip and the squared curvature of the whole body of the MT. In Fig. 4C, the MT started upward from a lower location. As a result, the MT grew toward the vertex and twisted around the vertex, avoiding the highest curvatures there. Eventually, the MT snapped to a downward configuration in which the curvature was more favourable. The exact nature of this apparent bifurcation is an issue for future investigation.

**Figure 4.**
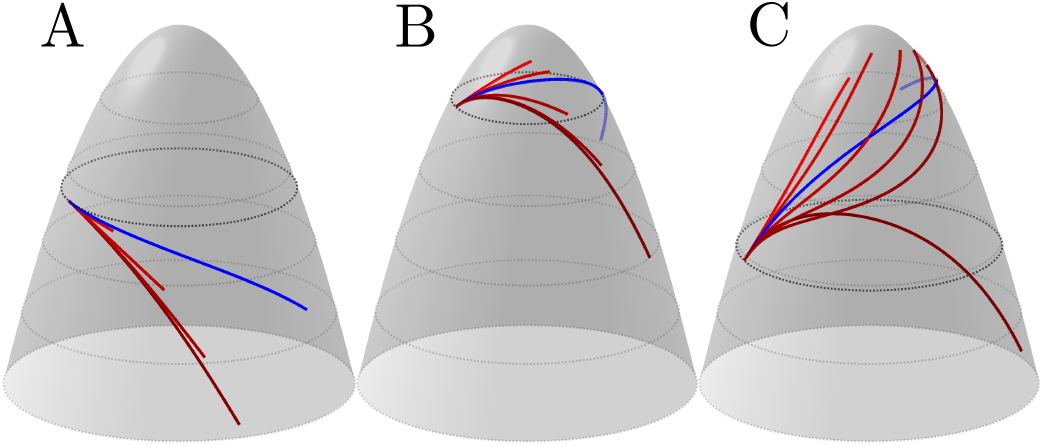
Elastic curves on a paraboloid of revolution. The surface parametrization is described in Appendix 5.2. Circles of latitude in light grey are at the same height in all figures, with a vertical spacing of 1 unit between each circle. The highest circle of latitude has a radius of 1 unit. Dark circles of latitude correspond to the starting heights of the curves in each figure. For each initial position and angle, a comparison group (similar to Fig. 3) of MTs was plotted. A: MTs starting at a height of 2.25 below the paraboloid vertex with an initial angle tangential to a circle of latitude, with lengths 1, 2, 3, 4. B: MTs with an initial starting location closer to the vertex with lengths 1.3, 1.5, 1.75, 2.5, 4. C: MTs with an initial starting location further away from the vertex with lengths 3, 3.5, 4, 4.5, 5, 5.5. MT lengths are chosen to show visually distinct shapes.

Geodesics in this scenario did not have particular significance in the context of curvature minimization; geodesics took on trajectories that persist in their tangential direction, regardless of whether this led to higher or lower curvature along the surface. This illustrated, again, how geodesics ignore the curvature of the surface. Note how the geodesics were not good approximations of the elastic curves except in the limit of short MT tips (corresponding to a high anchoring frequency).

### 2.4 Elastic Curves Exhibit Complex Shapes on Ellipsoids

Ellipsoidal geometries are especially relevant to recent protoplast experiments on MT orientation, where it has been suggested that geometric cues of this shape should bias MTs to be parallel to the long axis as the default configuration [12, 14]. We specialized to the case of spheroids; surfaces resulting from rotating an ellipse. We revisited this idea by examining the shapes of individual MTs in spheroids as MTs grow. In this case, the nature of MT-membrane anchoring — whether anchors fix the MT position and initial angle — was less clear in the absence of a rigid cell wall but experimental observations suggest that CMTs at least remain attached to the membrane [48, 6]. Similarly, there was a question of the stiffness of the membrane relative to that of the MTs. For simplicity, we kept the common assumption of a rigid surface where CMTs are fixed in position and angle by the anchors.

We started with a prolate spheroid, which has regions of higher curvature at the poles (top/bottom). Due to the symmetry of the spheroid, it sufficed to examine MTs anchored at a point anywhere along a single line of longitude in one hemisphere. For a MT closer to the poles, its shape was similar to those found in the paraboloid scenario until it was long enough to reach the region near the equator. Compare Fig. 5A and Fig. 4B — the MT initially deflected toward the pole at short lengths, then deflected away at larger lengths. When it deflected away, it was difficult to say whether the MT was more “longitudinal” vs. “transverse” due to the non-monotonicity in height and orientation along its body. As the MT grew longer and approached the opposite pole in Fig. 5B, there were competing influences from non-local sensing of curvature from both poles. The upper portion of the MT twisted toward the top pole, despite the energy cost, to avoid the bottom pole in a manner similar to what was observed from MTs on the paraboloid far from the pole, seen in Fig. 4C. Thus, as the MT lengthened, its shape alternated between attraction to and repulsion from the poles. Examples of elastic curves displaying similar behaviour with different initial conditions are shown in Appendix Fig. 10.

**Figure 5.**
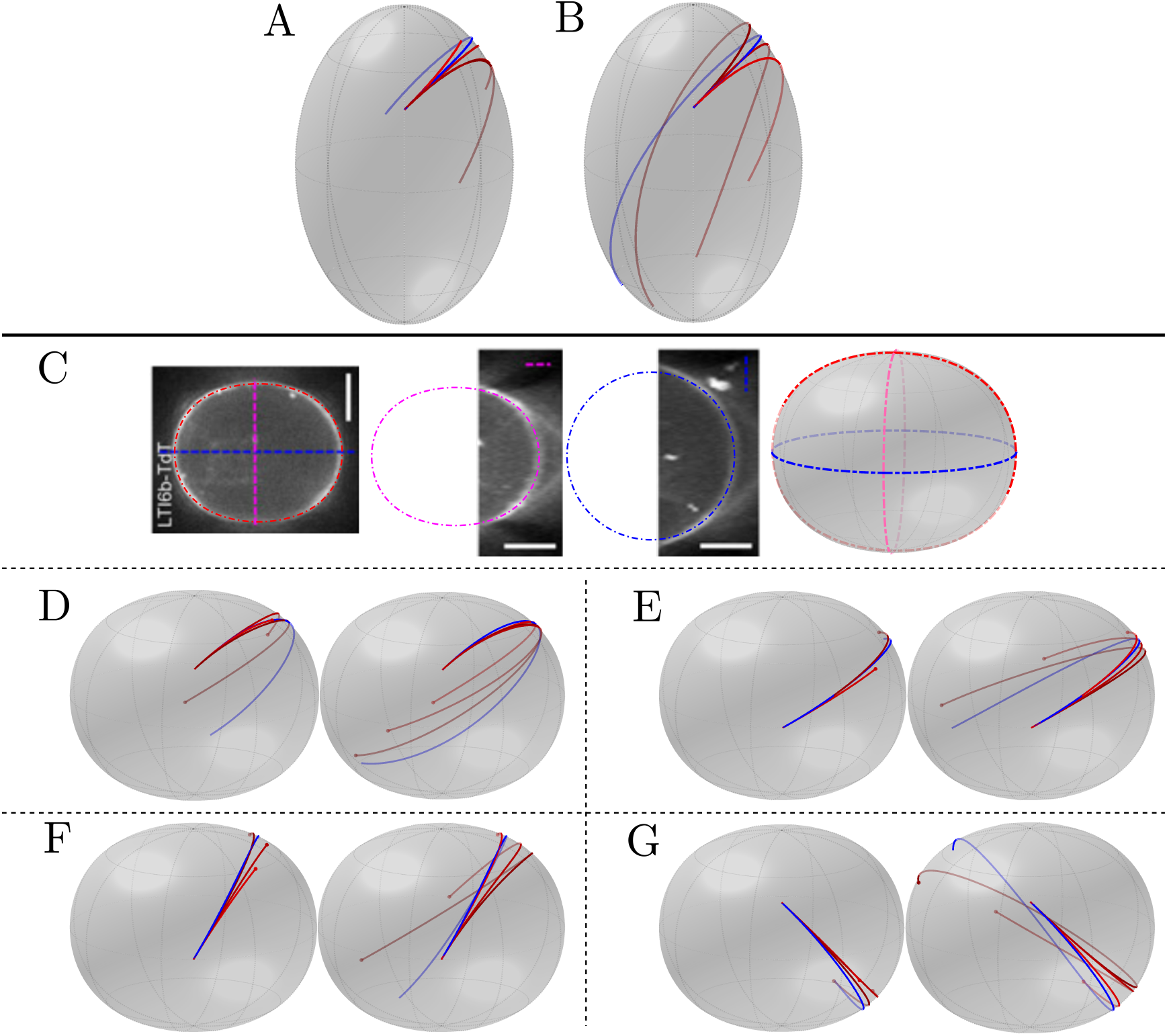
Elastic curves on spheroid-like surfaces of revolution (parametrizations described in Appendix 5.3) of various aspect ratios. A&B: prolate spheroids where the semi major axis is 1.5, semi minor axis is 1. MT comparison groups are shown in the same way as Fig. 3. The lengths of the MTs are {1, 1.5, 2, 3} (panel A) and {3, 4, 5} (panel B). C: oblate flattened spheroids with the same proportions as protoplasts in 15 *×*20 *µ*m microwells from [12] and the horizontal semi-axis scaled to have a length of 1 unit. Microscopy image taken from [12, Fig. 2E], with the red, pink, and blue dotted curves representing the cross sections of the fitted surface (right-most shape). D-G: MT comparison groups of elastic curves and geodesics on the spheroid in C. Within each panel, the longest elastic curve on the left figure serves as the shortest in the right figure. Lengths in each shape from left to right are as follows. D: {1, 2, 3} and {3, 3.5, 4 }. E: {1, 2} and {2, 3, 4}. F: {1, 1.5, 2} and {2, 3, 4}. G: {1, 2, 2.5} and {2.5, 3.5, 5}. Longer MTs for each panel are shown in Appendix Fig. 12 and 11.

Next, we considered oblate (flattened) spheroids fitted to microscopy images of confined protoplasts found in [12] (methods described in Appendix 5.3). It has been previously suggested that CMTs should orient parallel to the equator due to the lower curvature in this direction as compared to the higher curvature perpendicular to the equator [12]. In the same study, Colin et al. [12] found CMTs oriented perpendicular to the equator upon increasing protoplast pressure, attributing the bias against the default curvature cue to CMTs aligning to the direction with maximal tensile stress (corresponding toof cells packed in tissue such as s the direction perpendicular to the equatorial circle).

We examined two geometries, where the protoplasts were confined in microwells of varying aspect ratio: 15 *×*20*µ*m and 12 *×*40*µ*m. Because the two geometries produced qualitatively similar results, we displayed the fitted protoplast in the 15 *×*20*µ*m microwell case in Figs. 5C-G, with the 12 *×*40*µ*m case in Appendix Fig. 12. We varied the initial position and angle in this fixed geometry. In each of Figs. 5D-G, the MT comparison groups are presented for short (left spheroid) and longer MT lengths (right spheroid). Generally, shorter MTs deflected slightly toward the equator relative to the geodesic (e.g., shorter curves in Fig. 5F). For a narrow interval of intermediate lengths, the MT shifted toward the pole and for lengths above that (e.g., Fig. 5F right), it encountered the higher curvature regions along the equator, and shifted toward being parallel with the equator, rather than traversing it at a steep angle. For yet larger lengths, shown in Appendix Figs. 11 and 12, the MT began to sense the flat region of the opposing pole, causing the MT to shift toward the pole. This resulted in alternating movements similar to what was observed in the prolate scenario. Examples of elastic curves displaying similar behaviour with different initial conditions were shown in Fig. 5 panels D, E, and G, and Appendix Figs. 11 and 12. Together with the observations of MTs on paraboloids, this showed that the default geometric cues resulting from heterogeneous curvature and non-locality of MTs led to a variety of curves, rather than simply longitudinal or transverse curves. Our observation of MTs of intermediate length deflecting toward the poles within protoplast shapes while short and longer ones deflected the other way was particularly intriguing when considering the experimental observation [12] of transient alignment of CMTs toward the short axis in protoplasts under pressure.

## 3 Discussion

The recent interest in studying MT response to geometric cues has led to numerous studies, *in vitro* and *in silico*. There is increasing evidence of yet-to-be-elucidated active biological mechanisms to explain the observed emergent orientations of MT arrays. To understand the active processes involved, it is necessary to understand the passive processes that the biological system must regulate: the mechanics of MT polymers. Until now, mechanically realistic CMT models have been out of reach for complicated geometries. We have overcome this mathematical hurdle and numerically calculated these elastic curves. The resulting shapes revealed a general phenomenon: MTs sense curvature non-locally along their entire free tip length. Understanding this non-locality has provided several insights into the response of MTs to geometric cues, and on how plant cells may control the extent of curvature sensing.

Considering cylindrical geometries with edges (representative of cells packed in tissue such as those in Fig. 1A(i)) we found that edges cause MTs to deflect substantially from geodesics — even if the MTs start on flat faces away from the edges. When MT free tips are short relative to the radii curvature, curvature is sensed locally (flat in this case), and the MTs do not deflect significantly. When a longer free tip encounters the edges, the MT senses the large curvature at the edges, causing the MT *as a whole* to deflect. The cylinders shown in Fig. 3, when dimensionalized to have a half-width of 5*µ*m, have the corresponding MTs with tip lengths between 2.5-15*µ*m — within range of experimentally observed cell dimensions and

MT tip lengths [1]. Whereas previous models have examined the role of CLASP localization to regions of high curvature to stabilize MTs [4, 10], none has explained how MTs are able to persist in their direction around an edge. Anchoring proteins may be increased in concentration at edges where deflection is greatest, such that the free-tip-length–to–radius-of-curvature ratio is reduced. Then, the MTs become geodesic-like. On the other hand, anchoring proteins are not necessary at high concentrations in flat regions. This is consistent with the observation of CLASP proteins (associated with increased anchoring) localizing to regions of high curvature within Arabidopsis root cells [2]. This idea elucidates the potential dual-purpose of CLASP localization to regions of high curvature: allowing MT passage across edges by both MT tip stabilization and increasing anchoring to preserve directional persistence across the edges. In addition, other proteins such as MAP65 [13, 17, 31] may play a role in creating persistent transfacial bundles (Fig. 1A(i)) that act as guides for MTs to bundle with, assisting with overcoming sharp edges.

In geometries with competing influences of curvature, such as a rounded protrusion, the nonlocality of curvature sensing resulted in diverse MT shapes. For a MT initially oriented to the tip, it may first grow toward the protrusion tip since this direction results in less curvature than growing around the protrusion. However, as the MT reaches close to the protrusion tip, the high curvature causes the MT to snap back downward. For an initially transverse MT, there is a preference to bend away from from the protrusion. This is potentially a contributing factor to the observation of MT-depleted regions at the tips of trichome (Fig. 1A(ii)): through active processes (see, e.g., [44]), an initial transverse MT ring is created. New MTs, predominantly nucleating at shallow or (anti)parallel angles to transverse MTs, deflect away from the tip.

A similar phenomenon was seen in the case of ellipsoidal geometries, serving as models for protoplasts (Figs. 1A(iii) and 5C). Once a MT’s length spans the entirety of the cell, it exhibits alternating movement as it minimizes curvature by shifting into accessible low curvature regions and away from high curvature regions. This observation has potential implications for a recent protoplast study where, upon increasing osmotic pressure in protoplasts, MT orientation is observed to transiently bias parallel to the short axis (toward the pole of an oblate ellipsoid) before returning to be aligned with the long axis [12]. We speculate: could the default mechanics of MTs observed here have contributed to the behaviour seen in the tension study? Regardless, this work shows that, due to the extended nature of MTs, the nonlocal curvature sensing results in behaviours previously not considered.

In the case of a cylindrical cell, deflection was always longitudinal [45] making it more straightforward to conjecture about the influence of bending-energy minimization on CMT array orientation. With the more variable shapes addressed here, such a prediction is complicated by the complex curvature landscapes that induce non-monotone transformations of MT shape as a function of the free tip length. We leave our observations qualitative here and defer quantification of ordering to future studies that include MT interactions.

There are limitations associated with the assumption that CMTs are fully constrained to the surface. In the case of MTs encountering edges of sufficiently high curvature, it may be possible or even necessary for the MTs to lift from the membrane to bridge the edge. The curvature energy becomes extremely large for a MT that is fully constrained to a sharp edge. By pulling away from the membrane, a MT can round the same edge with a lower curvature thereby decreasing the energy. Thus, it may not be appropriate to directly apply this model to sufficiently sharp edges whose radii of curvature are less than the average anchor distance.

There is the related question of the contribution of thermal fluctuations. The rigidity of MTs as measured in vitro suggests that thermal fluctuations should not cause substantial deviation from the elastic curves we calculate in the present work. However, fluorescence imaging shows apparently softer MTs in the presence of MAPs [40]. This suggests that one strategy to control MT shapes may be through modifying their tendency to fluctuate, as examined in a previous model [25] in the context of protoxylem formation. Similarly, the apparent structural properties of MTs may be altered by the forces acting upon the filaments [34]. Future studies of the role of MAPs and cellulose synthase complexes in modifying the rigidity and shape of MTs [30] as well as including thermal considerations in our elastic curve model will be critical to reconcile these observations.

In addition, as mentioned previously, there is the question of how MTs are anchored to membranes in protoplasts and whether the membrane is deformable by the MTs when under osmotic pressure. Most existing CMT models treat CMTs as fixed on the membrane, even for protoplasts, which lack a rigid cell wall. The model of Mirabet et al. allows CMTs to detach from the membrane but treats them as geodesics (with stochastic fluctuations) while attached [35]. There remains the possibility in protoplasts of CMTs shifting laterally due to the absence of a cell wall, which presumably provides structural support for anchoring proteins. With this lateral freedom, the assumption of geodesic CMTs (corresponding to infinite anchoring) would be even less justified. Experimentally studying CMTs in isolation to characterize their shape has been challenging and our work may motivate future efforts to better understand the interplay between CMT-membrane interactions.

Despite these shortcomings, our work provides a foundation for understanding how geometric cues affect CMT organization. Modelling CMTs as geodesics is informative but limited in that it only reveals the impact that distances between points on a surface have on MT shape and organization and not the impact of curvature. The limitation is illustrated by the invariance of geodesic CMT models on surfaces that have a distance-preserving map (isometry) to a plane. Curvature does not have any consequence in such cases. The differences in geodesic MT arrays that may arise in cylindrical models are only due to cells having different sizes (e.g., height and width) and end-cap shapes. Such differences in shape lead to differences in distances between points, for example, for a MT traversing an end cap or wrapping around the perimeter of the cylinder. Cell size has been shown to have significant consequences on models of CMT organization [47, 10, 39, 46]. The invariance of geodesics on cylinders also applies to models of CMTs with finite persistence lengths, where the MTs are modelled as growing along geodesics with randomly sampled orientations [35, 14, 25]. This is because angles are preserved under isometries. A model that implements geodesic MTs (e.g., assuming high anchoring rates) on generalized cylinders of various cross-sections can simplify the computational effort required by taking advantage of the isometry and simulating on a plane. To study the mechanical consequences of curvature cues, it is necessary to study elastic curves. Our results provide intuition for the expected “default” MT shapes in various geometries. This can inform experimentalists on how to design studies (*in vitro* and *in vivo*) that disentangle active biological processes from the expected passive biophysical factors in array organization. For example, it may be beneficial to study systems such as large cylinders (e.g., *Nitella* internodal or Tobacco BY-2 cells) to eliminate the effects of MTs sensing non-local and competing curvature cues.

From a purely mathematical point of view, the method developed to find curvature minimizers and our analysis opens the possibility of studying the stability of elastic curves and the nature of bifurcations in various geometries. Understanding these bifurcations may also be of biological interest in other systems such as *Drosophila* organ development, as discussed in a recent study of compressive buckling of elastic curves on quadratic surfaces [53]. From a modelling point of view, the incorporation of this model into one of interacting MTs opens the possibility of more realistic CMT models in diverse plant cell geometries. This includes extending the results from this work on paraboloids and ellipsoids to more fully study trichome cells and protoplasts, respectively. Fig. 6 shows one possibility, where CMTs are modelled on a closed cubic prism with rounded edges resembling that of an *Arabidopsis* division zone root epidermal cell. This is in addition to geometries not studied here, such as pavement cells whose geometries can be modelled as a rounded prism (Fig. 6) with a cross-section similar to that of the second rightmost cylinder in Fig. 1B.

**Figure 6.**
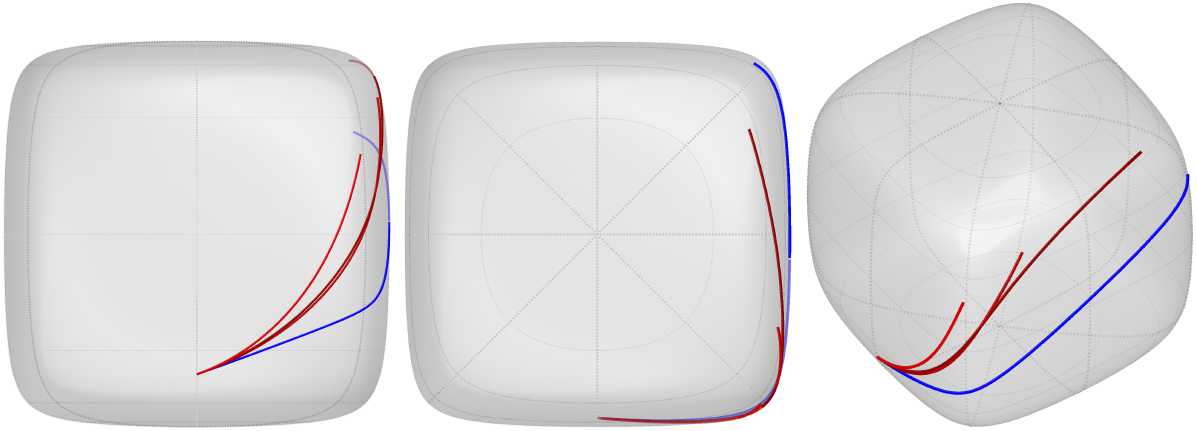
Cortical microtubules modelled as elastic curves on a cubic prism with rounded edges, resembling an Arabidopsis division zone root epidermal cell. The side length of the cube is 2, with elastic curves with the same initial position and angle of lengths 1.5, 2, and 3 representing the cortical microtubules. The blue curve is a geodesic with the same initial condition and a length of 3. From left to right: a direct view of one face of the prism, a top-down view (rotating the left-most view by 90^*°*^), and a perspective projection.

## 4 Methods

We studied a variety of shapes, and describe their mathematical parametrization by (*u, v*) *∈* ℝ^2^.The ℝ^3^ coordinates are thus three functions of (*u, v*), *x*(*u, v*), *y*(*u, v*), *z*(*u, v*), where the *x, y*- plane is the horizontal plane, and *z*-axis is vertical. To calculate the elastic curves in each case, we applied the numerical optimization method described in Sec. 4.2. Geodesics were calculated numerically (see, e.g., [9]) with standard ODE solvers (SciPy’s `scipy.integrate.ode` [51]).

### 4.1 Curvature Energy Functional

We want to calculate an energy-minimizing curve, ***γ***^*∗*^(*s*), representing a MT of length *L* constrained to a surface, *S*, and parametrized by arc length *s*. We define the energy functional,

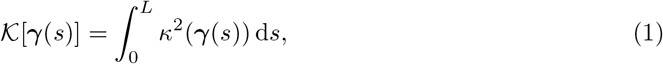

where, for each *s ∈* (0, *L*), we require 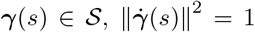, and 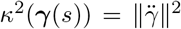, and we impose that ***γ***(0) = ***r***_0_ and 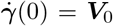, where ***r***_0_ is the position of the anchor end of the MT tip and ***V***_0_ is the unit tangent. Variables in ℝ^3^ are denoted with bold font and derivatives with respect to *s* use dot notation. The optimization problem of finding the minimizer of the energy functional, ***γ***^*∗*^(*s*), is solved using the numerical optimization method described next in Sec. 4.2.

At a high-level, there are two main approaches to finding the minimizer of the energy function ***γ***^*∗*^(*s*): optimize-then-discretize or discretize-then-optimize. The former leads to the Euler–Lagrange equations, which are differential equations that differ for each surface and are complicated to write down in general [36, 33, 53]. The differential equation must then be solved numerically. The latter alternative of first discretizing the curve is somewhat easier to get started with, allows the use of existing general-purpose optimization software, and is more readily adaptable to many surfaces; we describe this approach next.

### 4.2 Numerical Optimization: Problem Statement

For the purpose of numerically solving the optimization problem as well as ultimately simulating the dynamic multi-CMT interaction model, we pull the surface and MTs back into the plane. We define a *surface patch* as a mapping from a subset of the plane to *S*:

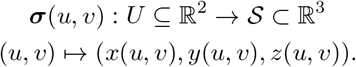

If the surface patch covers all of *S* then a curve on the surface can always be parametrized as

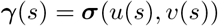

where we assume sufficient smoothness of ***σ***. We seek functions *u*^*∗*^(*s*) and *v*^*∗*^(*s*) that each map ℝ *→* ℝ, where *s ∈* [0, *L*] such that ***γ***^*∗*^(*s*) = (*u*^*∗*^(*s*), *v*^*∗*^(*s*)) minimizes the energy functional. Discretizing *s* into the set {*s*_*i*_ *}*for *i* = 0, …, *N*, we let *u*_*i*_ = *u*(*s*_*i*_) and similarly for *v*_*i*_. We chose

Chebyshev nodes of the second kind for the *s*_*i*_. Our search for *u*^*∗*^(*s*) and *v*^*∗*^(*s*) is now replaced by a search for *u*_*i*_ and *v*_*i*_. Let ***r***_*i*_ = *γ*(*s*_*i*_) = ***σ***(*u*_*i*_, *v*_*i*_) be the nodes along the curve. To keep the curve of a specified length, we have constraints on the spacing between nodes,

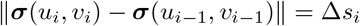

where Δ*s*_*i*_ = *s*_*i*_*− s*_*i−*1_. We discretize the energy functional (1) following the approach in [8], where elastic rods are discretized, and at each vertex, a discrete curvature is assigned,

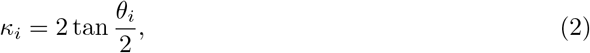

where *θ*_*i*_ is the turning angle between the corresponding displacement vectors ***e***^*i−*1^ and ***e***^*i*^, where ***e***^*j*^ = ***r***_*j*+1_ *−****r***_*j*_. The point-wise curvature is defined by averaging over the corresponding segment lengths 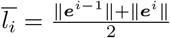. The curvature energy is given by the (discretized) integral,

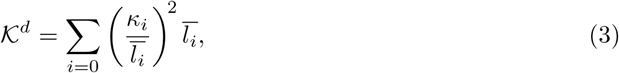

where the index starts from *i* = 0 due to the addition of a ghost point, explained below.

We are given the initial point ***r***_0_ = ***σ***(*u*_0_, *v*_0_) and implement the initial tangent condition by adding a ghost point ***r***_*−*1_ = ***r***_0_ *− h****V***_0_. The spacing between the initial and ghost points is chosen to be *s* = *h*_0_ *− h*_*−*1_. To make it more clear that the points ***r***_0_ and ***r***_*−*1_ are already determined and that the points *u*_*i*_ and *v*_*i*_ are to be determined, we write the functional (3) as 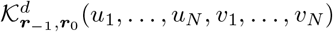. In all, we have the minimization problem,

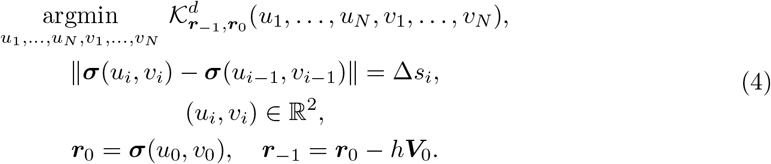

This is implemented directly into the trust-region optimizer from SciPy’s optimization library `scipy.minimize(method=‘trust region’)` [51]. Numerical tests are shown in Appendix 5.4.

### 4.3 Numerical Optimization: Practical Considerations

The optimization routine requires a suitable initial guess for the set of points (*u*_*i*_, *v*_*i*_). The initial guess is taken as either a line (in (*u, v*)) or a geodesic (of the surface).

To avoid the initial guess falling into a nonphysical minimum, the MT was “grown” to its final length. That is, final length *L* was sub-divided into sub-lengths, and the optimizer was used to iteratively solve on gradually longer lengths, using the previous solution (with a linear or geodesic extension) as the next initial guess.

## Acknowledgements

This work was supported by Natural Sciences and Engineering Research Council of Canada grants to E. Cytrynbaum, C. Macdonald, and G. Wasteneys. We thank participants of the Lorentz Workshop on “Modelling plant cortical microtubules” (2025) and the Banff International Research Station workshop on “Modelling Plant Microtubules” (2026) for insightful discussions on this work. We also thank Miranda Holmes-Cerfon and Keenan Crane for the helpful suggestions in approaching the optimization problem.

## 5 Appendix

### 5.1 Generalized cylinders

Generalized cylinders are surfaces obtained by translating a curve along a line. In this case, the curves are translated along a perpendicular line, giving upright cylinders. We assume the curve to be sufficiently smooth. For example, a circular cylinder is obtained by translating a circle along the axis perpendicular to the plane in which the circle lies.

#### 5.1.1 Squircular cylinder

The *squircle* is a family of closed curves that interpolates between a circle and a square [16]. The curve inscribed in the unit square can be parametrized using polar coordinates *θ∈* [0, 2*π*]

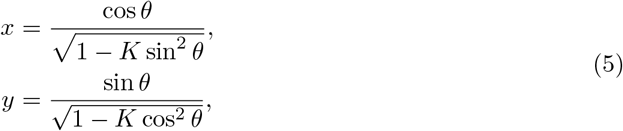

where the parameter *K ∈* [0, 1) characterizes how square-like the curve is, with the shape approaching a square as *K →* 1 (illustrated in Fig. 7).

**Figure 7.**
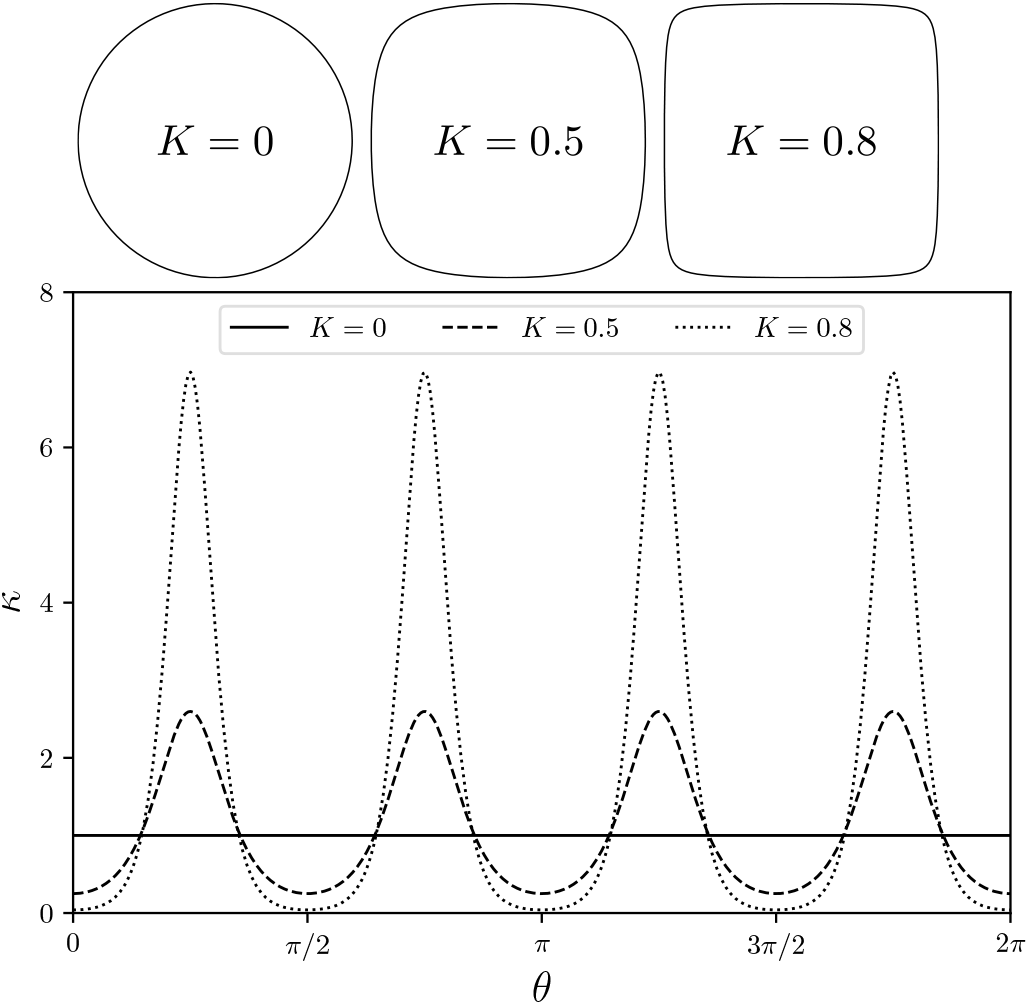
Shapes of squared-circles (*squircles*) parametrized from Eqs. 5. Top: the parameter *K* describes how square-like the shape is. Bottom: the curvature *κ* of the shape as a function of the polar angle *θ*.

The radius, as defined and measured from the centre to the right-most side of the shape (*θ* = 0), was 1 for all *K* values. The edges were located at odd integer multiples of *θ* = *π/*4, with increasing curvature at the edges as *K →* 1. As with any planar curve, the squircle can be extended into a generalized cylinder, which we refer to as the *squircular cylinder*, parametrized by coordinates (*u, v*) *∈* [0, 2*π*] *×* ℝ, with ***σ*** defined by
x

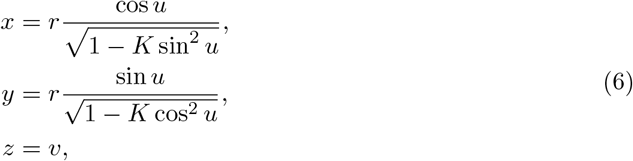

where *r* controls the “radius” of the cylinder. Fig. 7 and the top row of Fig. 3 use *r* = 1 (with the same cross-sections). The bottom row of Fig. 3 used *r* = 5 and *K* = 0.74. The squircular cylinder is not the solution of a physical packing problem, but we adopted it as a phenomenological model for the shape of a plant cell packed tightly among others like it.

### 5.2 Paraboloid of Revolution

The surface obtained by revolving the curve *z* = *−x*^2^ around the *z*-axis can be parametrized as

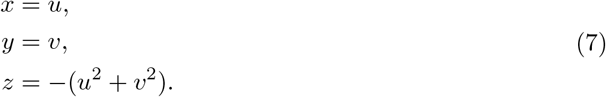

We used a Cartesian parametrization rather than a cylindrical one (i.e., in terms of height and azimuthal angle) to avoid potential issues with a singularity at the paraboloid vertex. The domain is ℝ^2^.

### 5.3 Spheroid

Our spheroid was a surface obtained by rotating the ellipse 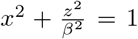 around the vertical axis. Here, 1 and the parameter *β* correspond to the major and minor axis respectively or vice versa depending on the relative magnitudes. The surface was parametrized using spherical coordinates,

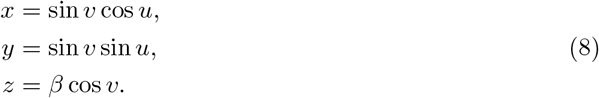

In this case, the domain was (*u, v*) *∈* [0, 2*π*] *×* (0, *π*). Setting *β* = 1.5 gives the prolate spheroid in Fig. 5A. Because the poles *v ∈ {*0, *π}* were singular points, the curves examined in this work were not close to the poles.

#### 5.3.1 Flattened Spheroid

The confined protoplast had its sides flattened due to being squished in the microwell. Similar to the squircle, it is possible to create a cross-sectional shape interpolating the ellipse and a rectangle with a parameter *K* describing its squareness [16],

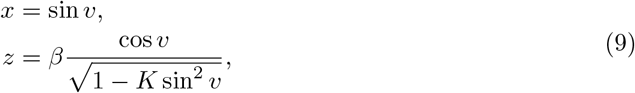

with the horizontal semi-axis of 1 unit and vertical semi-axis of *β* units. By visually fitting to the microscopy images in Fig. 1D and Fig. 2E in [12], we used *β* = 0.82 and *K* = 0.2 for the 15 *×*20*µ*m microwell and *β* = 0.78 and *K* = 0.3 for the 12 *×*40*µ*m microwell. These are shown in the dashed outline of each shape in Fig. 1(iii) and Fig. 8. These values were slightly different from the reported values of *β* = 0.88 and *β* = 0.86, respectively (Supplemental Material of [12]), where the authors fit the protoplasts to ellipses (*K* = 0).

**Figure 8.**
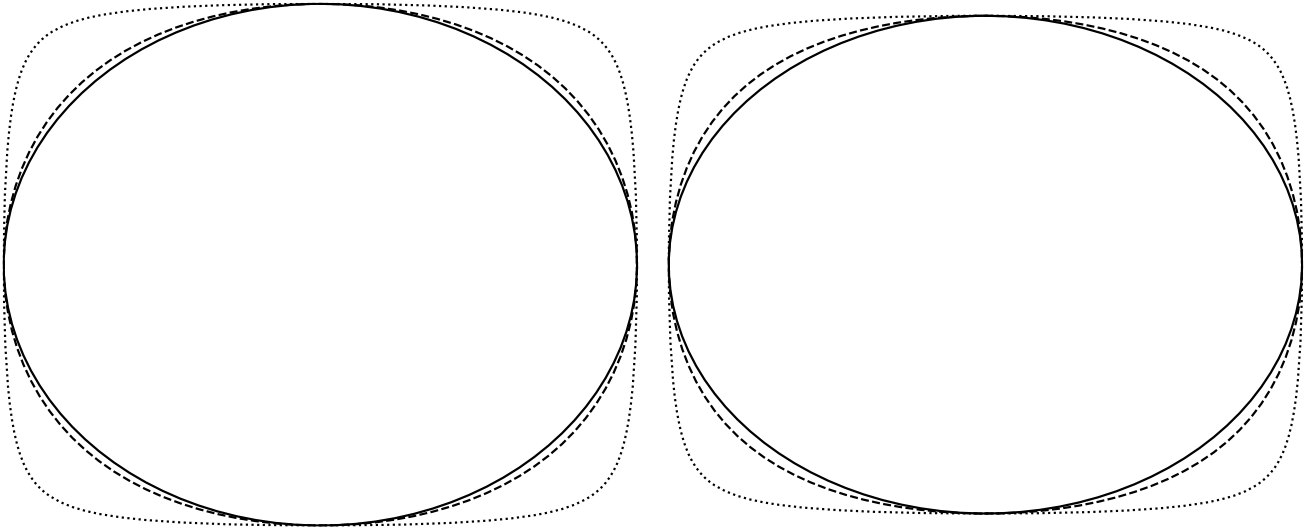
Shapes of squared-ellipses parametrized from Eqs. 9. Left: shapes with a horizontal semi-axis of 1 and vertical semi-axis of *β* = 0.82, corresponding to a scaled fit of the protoplast image in a 15*×* 20*µ*m microwell [12, Fig. 1D]. From inner-most to outer-most: *K* values (Eq. 9) of 0, 0.2, and 0.9. Right: shapes with a horizontal semi-axis of 1 and vertical semi-axis of *β* = 0.78, corresponding to a scaled fit of the protoplast image in a 12 *×*40*µ*m microwell [12, Fig. 2E]. From inner-most to outer-most: *K* values of 0, 0.3, and 0.9.

This shape was then rotated about the *y*-axis to give the parametrization,

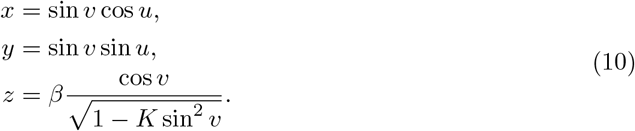

This gave the flattened spheroids in Fig. 5B and C.

### 5.4 Numerical Optimization: Numerical Tests

We tested the convergence of the optimizer on the cylinder by comparing the elastic curves calculated using the method presented here against numerically solving the Euler–Lagrange equations [5]. Let us denote the numerical solution of the Euler–Lagrange equations as *P* (*s*) = (*u*^*∗*^(*s*), *v*^*∗*^(*s*)), and the solution from numerical optimization as *p*(*s*_*i*_) = (*u*(*s*_*i*_), *v*(*s*_*i*_)). We measured the error as

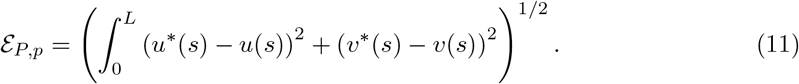

In the discrete case, it was given as,

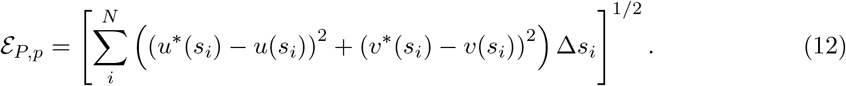

We started with the standard mapping for a unit cylinder, *S*_*c*_, given by

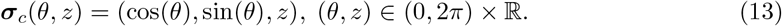

Here, *θ* represented the polar angle made between the position vector and the *x*-axis and *z* represents the height. The initial point used was ***σ***_*c*_(0, 0).

We then compared with an alternative mapping of a half-cylinder given by

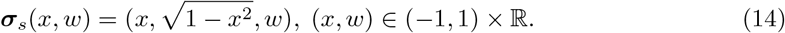

In this case, *x* represented the value along the *x*-axis with 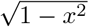 tracing the *y*-values of a semicircle, and *w* represents the height. The initial point used was 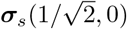 (equivalent to starting at (*π/*4, 0) in cylindrical coordinates). This was chosen to avoid the singularities at *u* = *±*1.

The numerical solution to the Euler–Lagrange equation found for the standard cylindrical coordinates denoted (*θ*^*∗*^(*s*), *z*^*∗*^(*s*)) on a fine mesh (node spacing smaller than 1*/*4000), and the points (*θ*^*∗*^(*s*_*i*_), *z*^*∗*^(*s*_*i*_)) were compared directly with the numerical optimization solutions (*θ*(*s*_*i*_), *z*(*s*_*i*_)) resulting from the standard cylindrical coordinates. To compare with the numerical optimization solution resulting from ***σ***_*s*_, we mapped the solution to the standard cylindrical coordinates (*x, w*) ↦ (arccos(*x*) *−π/*4, *w*).

The results for different initial angles were shown in Fig. 9. We found that the optimizer returns solutions with nearly identical accuracies between the surface mappings used; the filled and open markers, corresponding to the errors from each mapping, are nearly coincident. Through linear regression, the rate of convergence was found to be proportional to *N*^*−p*^, for *N* nodes, where *p* ranged from 2.2–2.3 among the different initial angles used.

**Figure 9.**
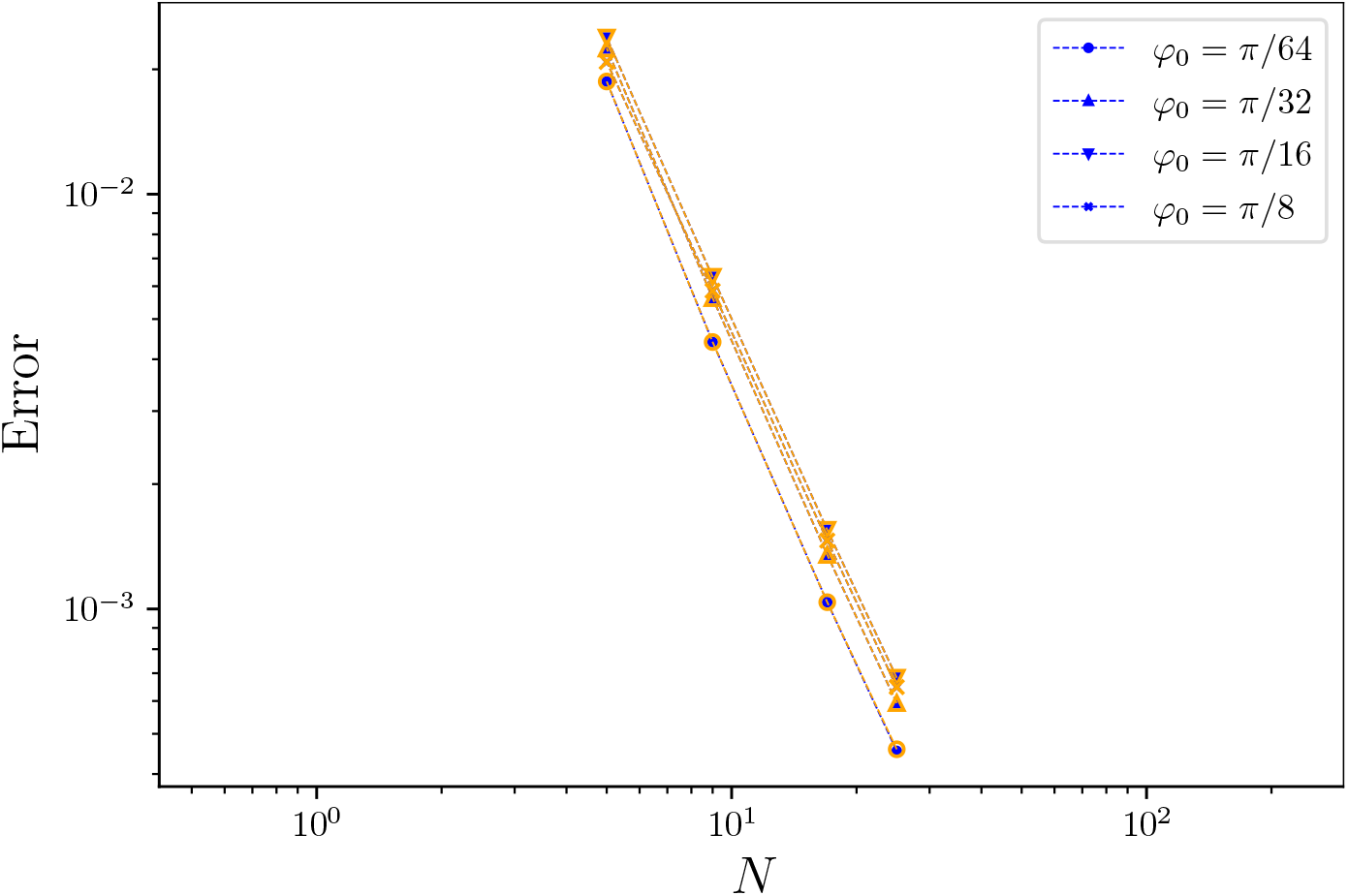
Convergence of the numerical optimizer applied to the discretized minimization problem Eq. 4 to the numerical solution of the corresponding Euler–Lagrange equation on for elastic curves on the circular cylinder. Errors are measured according to Eq. 12 as a function of the *N* number of points discretizing a curve of length 1. Error is plotted on a log-log plot, and *N∈* [5, 9, 17, 25]. Different initial angles, *φ*_0_, are represented by the different markers. The blue filled markers correspond to the solutions from the mapping Eq. 13, and the orange open markers are from the alternative mapping Eq. 14.

**Figure 10.**
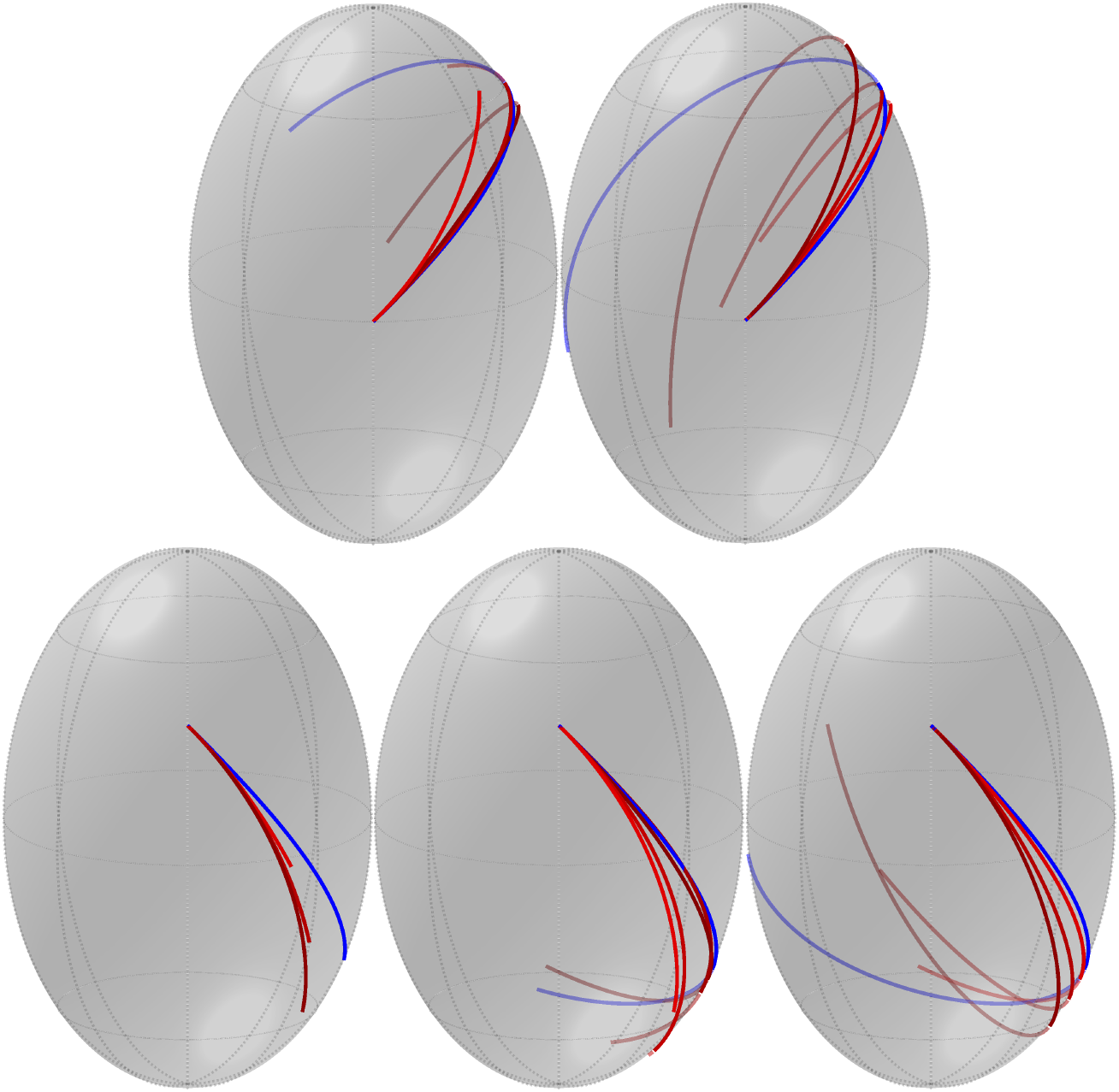
Examples of elastic curves (red) and geodesics (blue) on the same prolate spheroids as Fig. 5. Within each row, the curves start at the same position with the same initial angle. Progressing from left to right, the longest elastic curve on a given spheroid serves as the shortest elastic curve for the spheroid to its right. Geodesics in each spheroid have the same length as its longest elastic curve. Top row from left to right: MT lengths are *{*1.5, 2.5, 3.5*}*, and *{*3.5, 4, 5*}*. Bottom row: *{*1, 1.5, 2*}, {*2, 2.5, 3, 3.5*}*, and *{*3.5, 4, 5*}*.

**Figure 11.**
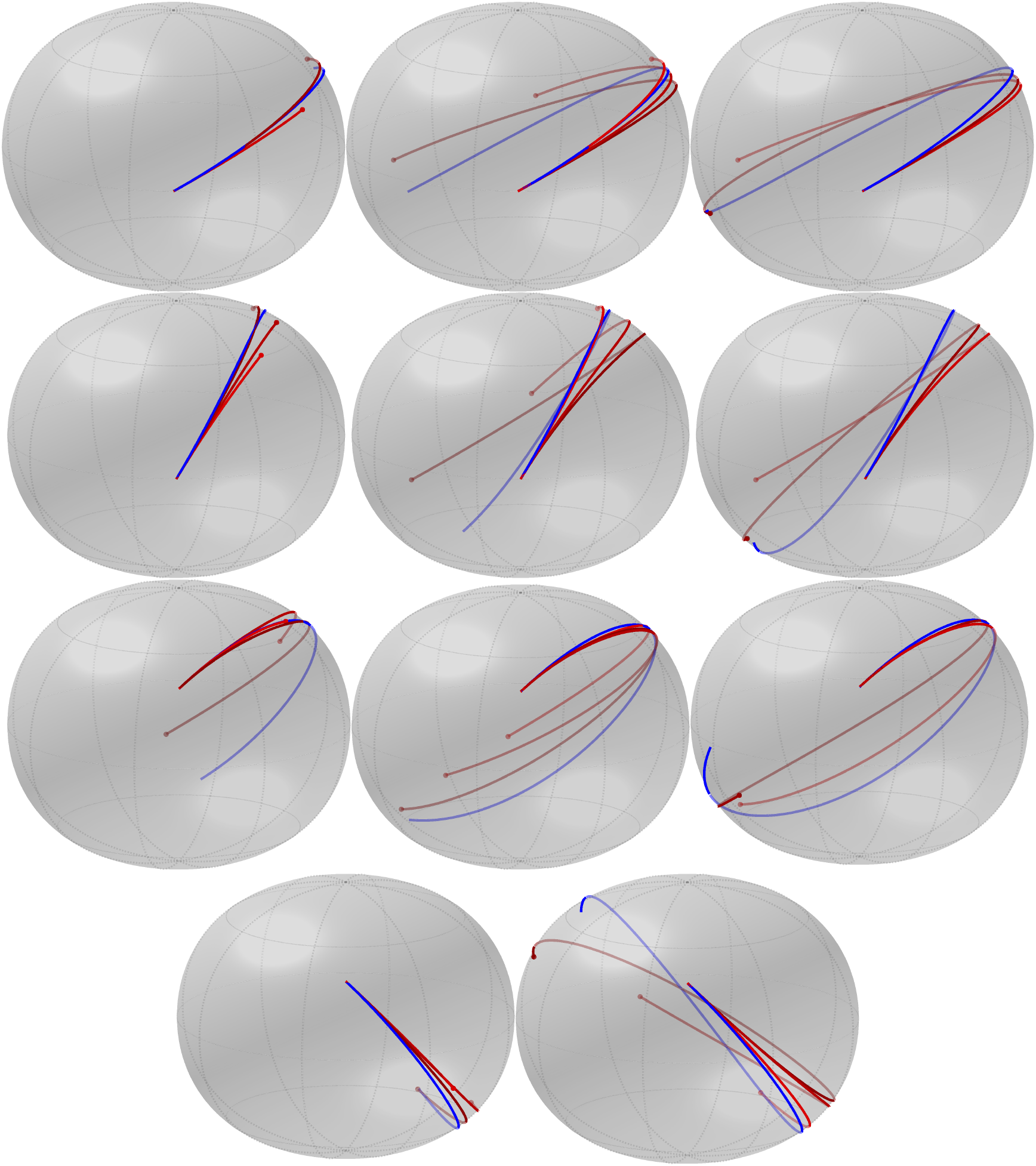
Examples of elastic curves (red) and geodesics (blue) on the same flattened oblate spheroids as Fig. 5D-G, corresponding the protoplasts in 15*×*20*µ*m microwells. Within each row, the curves start at the same position with the same initial angle. Progressing from left to right, the longest elastic curve on a given spheroid serves as the shortest elastic curve for the spheroid to its right. Geodesics in each spheroid have the same length as its longest elastic curve. The horizontal semi-axis is scaled to 1 unit. Top (first) row: MT lengths are *{*1, 2*}, {*2, 3, 4*}*, and *{*4, 5*}*. Second row: *{*1, 1.5, 2*}, {*2, 3, 4*}*, and *{*4, 5*}*. Third row: *{*1, 2, 3*}, {*3, 3.5, 4*}*, and *{*4, 5*}*. Fourth row: *{*1, 2, 2.5*}*, and *{*2.5, 3.5, 5*}*.

**Figure 12.**
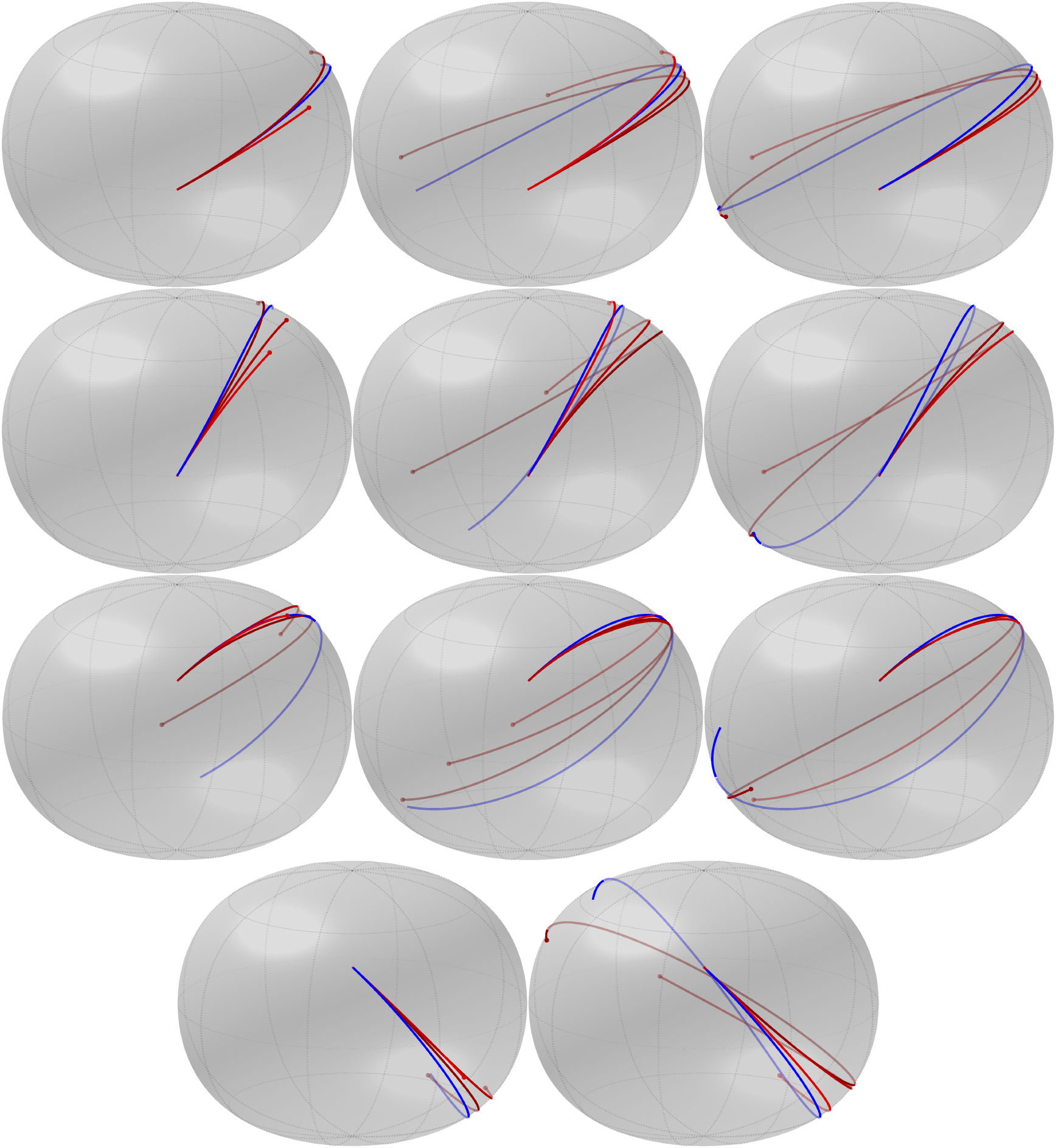
Examples of elastic curves (red) and geodesics (blue) on similar flattened oblate spheroids as Fig. 5D-G, corresponding the protoplasts in 12*×* 40*µ*m microwells. Within each row, the curves start at the same position with the same initial angle. Progressing from left to right, the longest elastic curve on a given spheroid serves as the shortest elastic curve for the spheroid to its right. Geodesics in each spheroid have the same length as its longest elastic curve. The horizontal semi-axis is scaled to 1 unit. Top (first) row: MT lengths are {1, 2}, {2, 3, 4}, and {4, 5}. Second row: {1, 1.5, 2}, {2, 3, 4}, and {4, 5}. Third row: {1, 2, 3}, {3, 3.5, 4}, and {4, 5}. Fourth row: {1, 2, 2.5}, and {2.5, 3.5, 5 }. The fit of this these shapes to the protoplast is shown in Fig. 1(iii).

The results for the numerical errors in Fig. 9 provided a general guideline for the number of nodes chosen for solving for the elastic curves in subsequent figures. The number nodes are displayed in Table 1.

**Table 1.** Number of nodes *N* used in each elastic curve calculation within each figure, in order of shortest to longest curves illustrated. Within each figure, the same number of nodes is used for the same lengths. Number of nodes in Fig. 5D-G are the same as those with the same corresponding initial conditions in Fig. 11. Similar for Fig. 12.

| | Number of nodes $N$ |
| --- | --- |
| Fig. 3A,B,C | {9, 16, 27, 36} |
| Fig. 3D | {30, 38, 45} |
| Fig. 4A | {15, 26, 35, 45} |
| Fig. 4B | {15, 18, 21, 23, 30, 35} |
| Fig. 4C | {35, 40, 45, 50, 54, 59} |
| Fig. 5A | {15, 21, 26, 35} |
| Fig. 5B | {35, 45, 54} |
| Fig. 10 first row | {21, 38}, {38, 53, 68}, {68, 83} |
| Fig. 10 second row | {21, 30, 38}, {38, 53, 68}, {68, 83} |
| Fig. 10 third row | {21, 38, 53}, {53, 60, 68}, {68, 83} |
| Fig. 10 fourth row | {21, 38, 45}, {45, 60, 83} |
| Fig. 6 | {42, 48, 54} |

### 5.5 Additional Plots

